# The Ensemble of Gene Regulatory Networks at Mutation-Selection Balance

**DOI:** 10.1101/2021.04.11.439376

**Authors:** Chia-Hung Yang, Samuel V. Scarpino

## Abstract

The evolution of diverse phenotypes both involves and is constrained by molecular interaction networks. When these networks influence patterns of expression, we refer to them as gene regulatory networks (GRNs). Here, we develop a quasi-species model of GRN evolution. With this model, we prove that–across a broad spectrum of viability and mutation functions–the dynamics converge to a stationary distribution over GRNs. Next, we show from first principles how the frequency of GRNs at equilibrium will be proportional to each GRN’s eigenvector centrality in the genotype network. Finally, we determine the structural characteristics of GRNs that are favored in response to a range of selective environments and mutational constraints. Our work connects GRN evolution to quasi-species models, and thus can provide a mechanistic explanation for the topology of GRNs experiencing various evolutionary forces.

## 1 Introduction

Molecular networks influence both macro- and micro-evolutionary processes [1, 2, 3, 4, 5]. But, how might they themselves evolve? A recent comparative study of regulatory networks found that their structures often exist at the edge of critically, straddling the border of chaotic and ordered states [6]. That biological regulatory networks should exhibit the kind of dynamic stability associated with near-critical networks has been theorized as adaptive, both from the perspective of functional robustness [7] and their ability to effectively process information [8]. However, there is also empirical and theoretical evidence for the importance of change in these networks, e.g., if species must evolve to meet shifting environmental or ecological selection pressures [9]. This tradeoff between robustness and evolability is hypothesized as an explanation for the common “small-world” property in biological networks [10]. Nevertheless, foundational work on self-organized criticality and 1/*f* noise demonstrated that dynamical systems embedded in a spatial dimension, e.g., biological regulatory networks, might naturally evolve to near-critical states [11, 12]. Therefore, one could observe near-critical networks in nature that are derived from constraints, as opposed to directly optimized by selective forces.

Focusing specifically on interactions that modulate expression, recent studies have hypothesized how various evolutionary forces shape the structure of gene regulatory networks (GRNs) [13, 14, 15, 16]. Analyses of transcription factors [17, 18], mRNA profiles [19] and comparative genomics [20] suggest that gene duplication/loss have a substantial contribution to divergent gene regulation. Moreover, several mathematical models of GRN evolution were introduced to encompass duplication events [21], selection on functional dynamics [22], horizontal gene transfer [23], correlated mutations on genomes [24], and non-genetic inheritance [25]. Force et al. [26] computationally showed that subfunction fission following duplication events can lead to a modular structure of GRNs. Similarly, Espinosa-Soto and Wagner [27] demonstrated that sequential adaptation to newly specialized gene activity patterns can increase the modularity of GRNs. Conversely, GRNs are hypothesized to emerge largely as a by-product of the progression towards some optimal state, via some combination of negative-feedback regulation [28], the rate of molecular evolution [29], tradeoffs between robustness and evolvability [6], and self-organization of functional activity [30].

In principle, existing frameworks that model evolutionary dynamics can be applied to the evolution of GRNs. Ideally, and hypothetically given “omniscience” over the genomes—including comprehension of every fundamental interaction between molecules—one can reconstruct inter-dependencies among genes and obtain GRNs from a bottom-up approach. Of course, this ambition is far from practical and even sounds like a fantasy. Yet, it shows that GRNs are essentially a direct abstraction of the genotypes. This abstraction is not only central to the omnigenic perspective of complex traits [31], but it also motivates a theoretical framework of regulatory circuit evolution [32]. Over the past two decades, several models of GRN evolution have been proposed [33, 34, 35, 36, 37]. The resulting models have influenced our understanding of diverse phenomena including canalization [38, 34], allopatric speciation [39, 36, 37], expression noise [40], and the structural properties of GRNs themselves [41, 27, 35].

However, to date, the existing models of GRN evolution remain largely computational, and the complexity of genetic interactions impedes more advanced theoretical analyses of these models. Our ambition in this work is to derive analytical conclusions for models of GRN evolution, with aids from known theoretical establishments in population genetics and quasi-species theory [42]. Population genetics describes how the frequency of different alleles change over time in a population mechanistically through evolutionary forces [43], in which mathematical models usually focus on a finite-sized population and a few loci. Quasi-species theory, on the other hand, concentrates on the balance between selection and mutation in an infinitely large population, where genotypes with a higher dimensionality are incorporated [44]. Literature has provided exact solutions for the steady distribution of genotypes along with their global convergence in quasi-species theory [45, 46, 47] under the assumption of irreducible and primitive transition matrices. This assumption was latter proposed to correspond to the mutational accessibility among genotypes with non-zero fitnesses [48]. It is thus not hard to vision that, when extended with complex genetic interactions, the stationary solution of a population-genetic or quasi-species model implicates the balanced distribution of plausible GRNs under the focal evolutionary forces. Nevertheless, it remains to be shown that such assumptions are valid for high dimensional genotype-phenotype maps associated with gene regulatory networks.

Here, we develop a quasi-species model describing how the structure of GRNs are shaped by a combination of selection and mutation. First, using this model we study the dynamics of GRN evolution in an infinitely large population with non-overlapping generations in a constant environment. By depicting the mapping between genotypes and phenotypes through the GRNs [37], we mechanistically recover the key assumption in the literature mentioned above and prove that the dynamics always converge to a stationary distribution over GRNs. Then, assuming binary viability, identical reproductivity, and rare mutation, we analytically show that the frequency of GRNs at mutation-selection balance is proportional to each GRN’s eigenvector centrality in a sub-graph of the genotype network [49, 50, 51, 52]. Finally, we determine the structural motifs associated with GRNs that are favored in response to a wide variety of selective regimes and regulatory constraints. We discuss the implications of our results in the context of the evolution of complex phenotypes and the challenges of studying GRN evolution.

## 2 Models

### 2.1 Quasi-Species Model with Selection, Reproduction, and Mutation

We begin with a quasi-species model that incorporates selection, reproduction, and mutation: The viable individuals in the current generation reproduce and generate their offspring, which may possibly mutate, and undergo selection to form the next generation. This phenomenological modeling scheme has frequently appeared in existing literature for a deterministic dynamics or a stochastic Markovian process perspective [44, 45, 46, 48, 47]. Yet, we shall see shortly that basic probability theory assists us to construct the model in a bottom-up fashion and leads to a probabilistic interpretation of various parameters. We additionally impose a few assumptions to the model, including a.) an infinitely large population size, b.) non-overlapping generations, c.) asexual reproduction, d.) a constant reproductivity of each genotype and a fixed selective environment over time, and e.) that any single-locus mutation has a non-zero chance to occur per generation.

Suppose that *I*_*t*_ represents an individual randomly sampled from the population at generation *t*. Let *g* (*I*_*t*_) and Ψ(*I*_*t*_) be its genotype and the event that *I*_*t*_ is viable respectively. We will further denote by *I*_*t*−1_ → *I*_*t*_ the event that the randomly sampled individual at generation *t* − 1, namely *I*_*t*−1_, reproduced and generated the randomly sampled individual at generation *t*, namely *I*_*t*_. We will also write 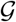 to represent the set of all plausible genotypes.

For any genotype 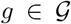, we are interested in its prevalence in the population at a given generation *t after* selection. In other words, we would like to know the probability that we observe a randomly sampled individual at generation *t* with the genotype *g*, given the fact that the sampled individual is viable. Applying the Bayes’ theorem, this focal conditional probability becomes

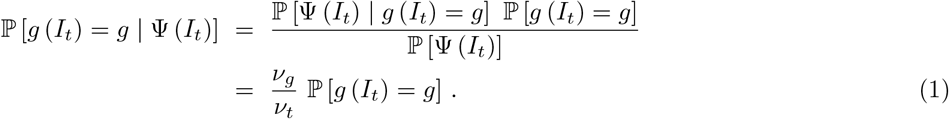

For simplicity we adapt the abbreviation *v*_*g*_ = ℙ [Ψ(*I*_*t*_) | *g* (*I*_*t*_) = *g*] and *v*_*t*_ = ℙ [Ψ(*I*_*t*_)], which are equivalently the survivial probability or the **viability** of genotype *g*, and the average viability at generation *t* respectively.

What we have left in equation (1) is the probability that a randomly sampled individual has genotype *g before* selection. The derivation of ℙ [*g* (*I*_*t*_) = *g*] relies on two observations: First, the genotype of individual *I*_*t*_ arose from mutation and the unique genotype of its parent; second, this parent individual must be viable. The event *g* (*I*_*t*_) = *g* is hence partitioned^1^ by the joint events 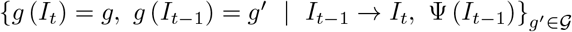. So we have

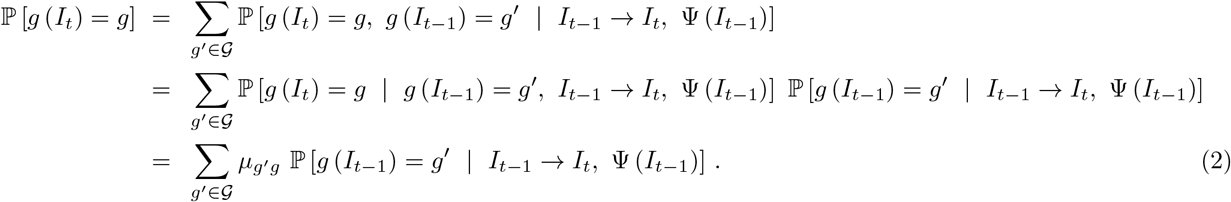

Again we abbreviate 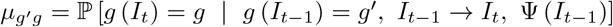 that shows the **mutation probability** from genotype *g*′ to genotype *g*.

It remains to resolve ℙ [*g* (*I*_*t*−1_) = *g*′ | *I*_*t*−1_ → *I*_*t*_, Ψ(*I*_*t*−1_)] in equation (2), which is the probability that the parent of a randomly sample individual at generation *t* has genotype *g*′. Applying the Bayes’ theorem once more, this probability becomes

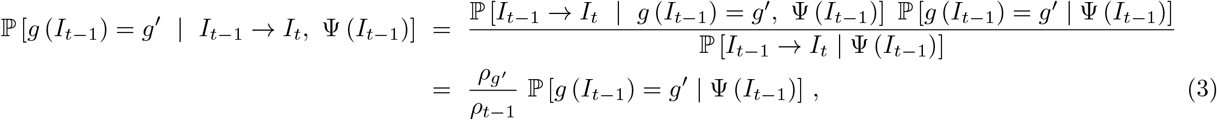

where 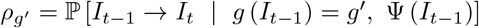 is the **reproductivity** of genotype *g*′, and *ρ*_*t*−1_ = ℙ [*I*_*t*−1_ → *I*_*t*−1_ | Ψ(*I*_*t*−1_)] is the average reproductivity at generation *t* − 1. Note that, instead of defining reproductivity as the number of offspring of an individual, the probabilistic formulation conversely describes, when sampling from the infinitely-sized next generation, how likely we will observe an offspring of the focal individual.

More importantly, we see that equation (3) leads us back to the focal conditional probability that, at generation *t* − 1, a randomly sampled individual has genotype *g*′ given that it is viable. Combining (1) to (3), we obtain the master equation for the simple quasi-species model that integrates selection, reproduction, and mutation of genotypes:

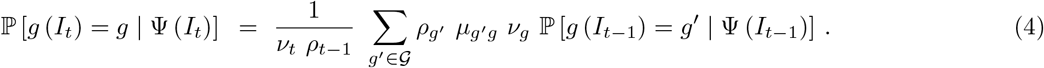

### 2.2 Pathway Framework of GRNs: Representing Genotypes by Expression Behavior

In existing literature of quasi-species theory, the model parameters are usually arbitrarily tunable or follow a particular distribution for simplicity. Hypothetically, these parameters depend on the resultant phenotypes of the genotypes, and any genotype-phenotype mapping reflects constraints and provides information of the model parameters. Our previous work has proposed a modeling approach, termed the **pathway framework**, to describe how the structure of GRNs varies due to genetic changes and how they respond to a given selective pressure [37] (which we summarize below; see its formal mathematical formulation in Appendix A). In the current work, we apply the pathway framework of GRNs to the quasi-species model (4) as a simple genotype-phenotype mapping for parameterization.

We note that the pathway framework is not without a handful of presumptions and thus restricts to rather specific GRNs. Nevertheless, the pathway framework GRNs play the role of an informative genotype-phenotype mapping which evokes some mechanistically interpretable parameterization for models in quasi-species theory. As a teaser, we shall in later sections that this naive and particular genotype-phenotype mapping through gene regulation surpasses the key assumption in existing quasi-species theory when proving the global convergence to a stationary solution. The pathway framework may seems an arbitrary and perhaps oversimplified choice to encapsulate the genotype-phenotype mapping; future works can indeed incorporate more realistic modeling frameworks to GRNs to strengthen the conclusion of global convergence.

The key of the pathway framework is to conceptualize alleles of genes as “black boxes” that encapsulate their expression behavior. Expression of a gene is triggered by some protein called the transcription factor, which is followed by a series of procedures to synthesize the protein product. The pathway framework of GRNs extracts the allele of a gene through this input-output relation, i.e., the activator protein(s) of gene expression and the protein(s) it produces. Regulation between two genes naturally arises once one gene’s protein product involves in the activation of the other’s expression (see Figure 1). Furthermore, these input-output relations of gene expression serve as the “inherited” reactions through which external environmental stimuli and internal chemical signals of proteins propagate to develop the phenotype. The pathway framework hence represent the genotype as the input-output relation of each gene’s expression behavior, where the corresponding GRN is constructed accordingly, and it considers the collective state of proteins as the resulting phenotype.

**Figure 1:**
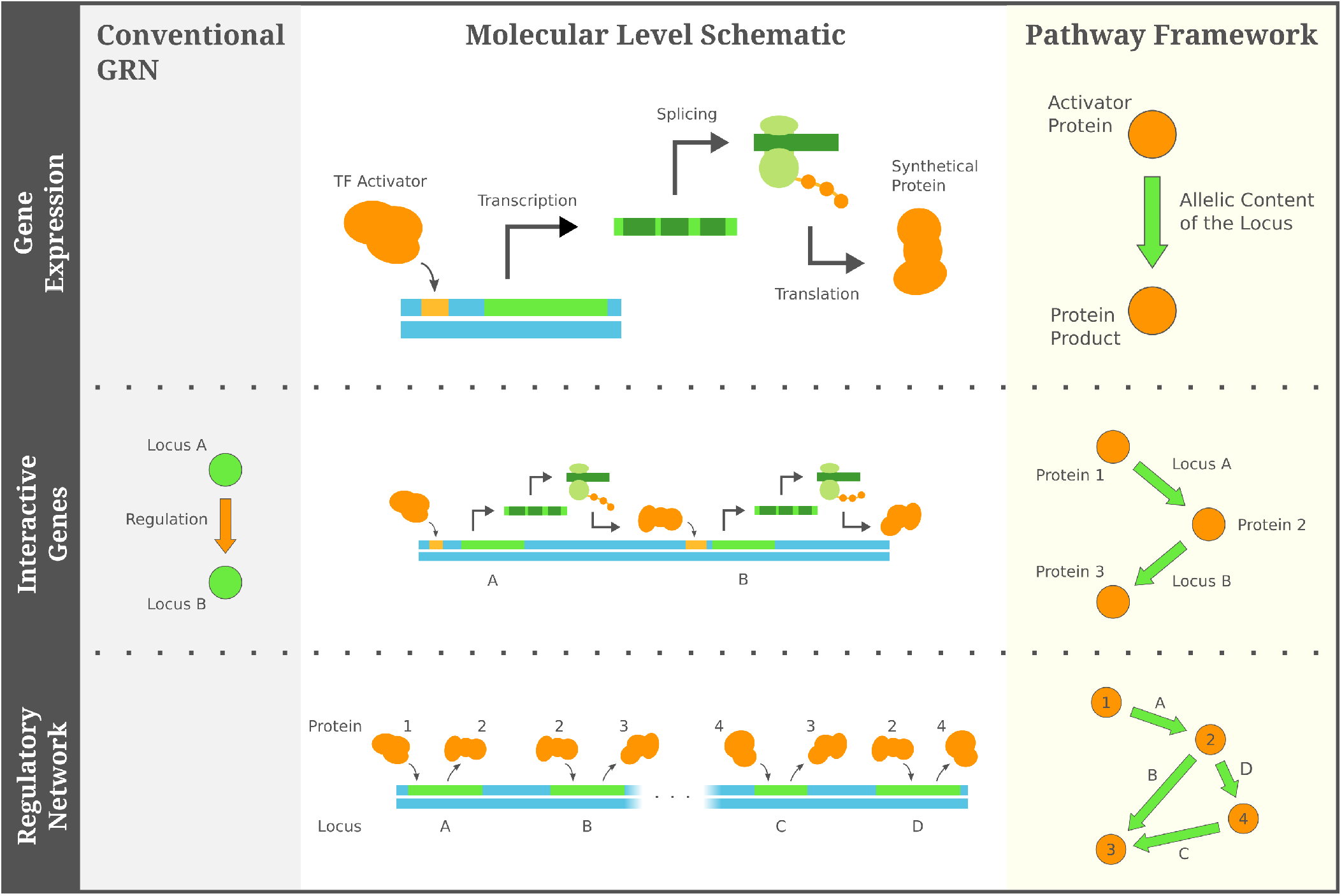
The pathway framework interprets a GRN as an abstraction of the expression behavior of the genotype. In this framework a GRN consists of edges indicating the input-output pair of a gene’s expression, from which transcriptional regulation between genes can be recovered, and it is arguably a more compact representation than the conventional notion of GRNs.

In this work, we focus on a minimal pathway framework of GRNs which integrates a few additional assumptions: First, we presume there is a constant collection of proteins Ω that can *possibly* appear in the organisms, and the state of a protein is binary, which indicates whether the protein is present or absent in an organism. Second, assuming that any gene’s expression is activated a single protein and produces a single protein product, the allele of the gene becomes the ordered pair of protein activator/product. If the protein activator is in the present state, the allele of the gene turns the state of the protein product to presence. Third, there is a fixed collection of genes Γ in the organisms, and the allele of each gene can be any pair of activator/product in the constant collection of proteins. Forth, the external environmental stimuli, if any, specify some activator proteins in the constant collection Ω and turn their state to presence.

Under these assumptions, a GRN can be transformed from its conventional notion, where nodes in the network represent genes and the edges shows regulation among them, into a more compact format such that the nodes are exactly the constant collection of proteins and the directed edges describe the expression behavior of alleles of genes (see Figure 1). Hereafter, if not otherwise specified, we refer the term GRNs to those in the compact format under the pathway framework, yet it is noteworthy that the two constructions are merely different representations of the expression behavior of the same underlying genotype. While the set of all possible genotypes is denoted by 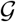 in section 2.1, we abuse the notation 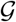 for their corresponding GRNs as well, and we write 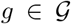 to refer to a possible genotype/GRN. Given the constant collection of proteins Ω and genes Γ, the set of all possible GRNs is determined: A possible GRN is a network among Ω with |Γ| directed edges, each of which is labeled by a gene in Γ and points from any protein activator to any protein product in Ω.

The pathway framework provides an approach to model evolutionary mechanisms, such as random mutation and natural selection, through graphical operations and structural characteristics on the GRNs. Mutation at a gene changes its allele stochastically, which is essentially a random process over all possible pairs of protein activator/ product in the constant collection Ω excluding the original allele. In the corresponding GRN, mutating the allele of a gene is equivalent to rewiring the directed edge labeled by the focal gene. On the other hand, selection is usually characterized as some phenotypic response against the environment. Specifically, since a phenotype is developed through the cascading of internal signal of protein appearances starting from the external environmental stimuli, the binary state of a protein in the resulting phenotype corresponds to its reachability from the stimulated proteins in the GRN. The viability and the reproductivity of a genotype can therefore be modeled as functions of node reachability in the GRN. For example, in the case study in section 3.2, we will consider a simple scenario where the mutation at each gene is independent and the outcome is uniform among all possible alleles, and that the viability is 1 if some phenotypic constraint is satisfied or 0 otherwise. We explore more complex scenarios in later sections.

### 2.3 Genotype Network: a Space of Mutational Relationship between GRNs

Previous literature has developed the concept of the genotype network, which captures how various genotypes transition from one to another through mutations (not necessarily just point mutations) and/or recombination [49, 53]. Here, we adopt the genotype network to describe the mutational connection between GRNs. The genotype network of GRNs is a undirected network of networks, where every possible GRN becomes a mega-node, and two mega-nodes are connected if the two corresponding GRNs only differ by the allele at a single locus. In other words, an edge between two mega-nodes in the genotype network represents a single-locus mutation between GRNs (Figure 2). Sometimes, instead of concentrating on all possible GRNs, we focus the mutational relationship between a subset of them. A particularly remarkable scenario is to constrain the GRNs on the binary state of proteins of the resulting phenotype. For instance, one common phenotypic constraint is to focus on the GRNs with equal fitness under selection, and the consequent induced subgraph of the genotype network is known as the neutral network [51, 53], which captures mutational transition between GRNs that are selectively neutral.

**Figure 2:**
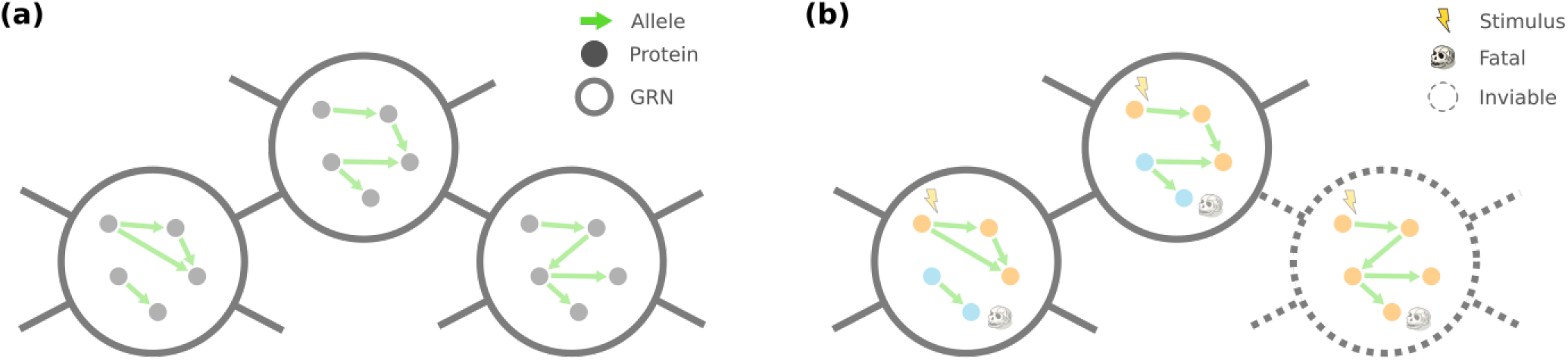
(a) Genotype network of GRNs, where, under the pathway framework, two mega-nodes (GRNs) are connected if and only if thy differ by one edge rewiring. (b) Neutral network of GRNs, where inviable mega-nodes are removed from the genotype network. In this illustrative example inviability is modeled as a regulatory pathway from the stimulus to the protein product with a fatal effect.

We emphasize two important properties of a genotype network of GRNs and its induced subgraphs under the pathway framework. First, because the underlying collections of proteins and genes are fixed, and a mutation at any gene can lead to a mutant allele that points from any protein activator to any protein product, each GRN has the same number of mutational neighbors. As a result, the genotye network of GRNs is in fact a regular graph. Second, for any phenotypic constraint, we show that the resulting induced subgraph of the genotype network is connected. In other words, there always exists a sequence of single-locus mutations between two GRNs such that the involved GRNs all satisfy the arbitrary given constraint on their phenotype. The guaranteed connectedness also applies to the neutral network of GRNs, where the phenotypic constraint corresponds to protein states leading to the same fitness.

We leave to Appendix C the detailed proof for the connectedness of a subgraph of the genotype network induced from arbitrary phenotypic constraint, and we only provide a brief outline here. The proof is based on a few observations of the pathway-framework GRNs. Under the presumption of binary protein state, there naturally exist some protein activator/product pairs that are “redundant” in terms of the resulting phenotype. Such redundancy manifests when the product is simply the activator itself, or when multiple genes the same activator/product pair.

Furthermore, given a phenotypic constraint, one can come up with a family of “naive” GRNs that satisfy the constraint. Specifically, such a “naive” GRN is constructed by (a) for each required-present protein, assign it as the product of a gene with an external stimulus as the activator, and (b) assign the rest of genes with redundant activator/product pairs. Our proof in Appendix C systematically finds a mutational trajectory between two GRNs *g*_*s*_ and *g*_*t*_ satisfying the phenotypic constraint. This trajectory consists of three segments — between *g*_*s*_ and a naive GRN *g*′, between *g*′ and another naive GRN *g*″, and finally between *g*″ and *g*_*t*_ — and all GRNs traversed by the mutational trajectory also satisfy the given phenotypic constraint.

## 3 Analyses

### 3.1 Convergence to a Stationary Distribution of GRNs

Our main result shows the convergence of the quasi-species model (4) under the pathway framework, and we derive the stationary distribution over possible GRNs. We begin with noting some groups of GRNs whose probability to be observed is relatively straightforward in the model. First, for any GRN *g* with a zero viability, i.e., *v*_*g*_ = ℙ [Ψ(*I*_*t*_) | *g* (*I*_*t*_) = *g*] = 0, the probability to observe *g* from a randomly sampled individual that has survived selection is also zero. Formally speaking, denoting those GRNs with a non-zero viability by 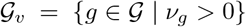, we have ℙ [*g*(*I*_*t*_) = *g* | Ψ(*I*_*t*_)] = 0 for each 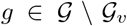 and at any time *t*. Second, denote the GRNs with a zero reproductivity by 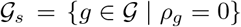. Since GRNs 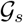 do not contribute to the offspring, their probability to be observed solely depends on the other GRNs 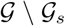. In particular, for each 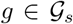 and at any time *t*, we have 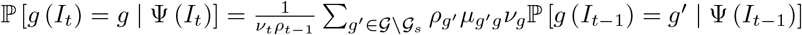.

It is thus useful to only keep track of the GRNs with a non-zero viability and a non-zero reproductivity. Hereafter, we consolidate the focal conditional probability for every 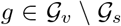 at gene ation *t* through a column vector **p**^(*t*)^. We write the *i*_*g*_-th entry of **p**^(*t*)^ as the one that corresponds to *g*, namely 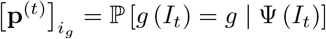. The master equation (4) can also be rewritten in a matrix format. Specifically, we denote by **T** a semi-transition matrix whose entry at the *i*_*g*_-th row and the 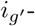th column is 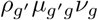, for any pair of 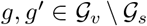. We in addition have another matrix **R** to capture the transition from 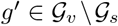 to 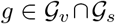, whose entry at the *i*_*g*_-th row and the 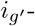-th column is again 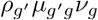. With these matrix notations, and along that the product of the average viability and the average reproductivity 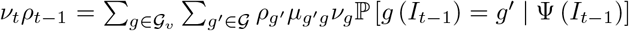 since 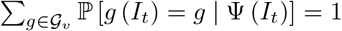, the master equation (4) therefore becomes

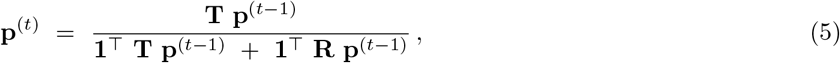

where we use the notation **1**^T^ for the row vector of ones with the proper length.

The matrix **T** plays a key role in the master equation (5), and it has a nice property that all its entries are positive. Since **T** corresponds to transition between GRNs 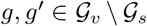, the relative reproductivity 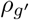 and the viability *v*_*g*_ are both positive. Next, we must show that the mutation probability 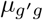 is positive as well. Recall that, when constructed through the pathway framework of GRNs, the subgraph of the genotype network induced by any phenotypic constraint is connected (see section 2.3 and Appendix C). More formally, the connectedness among GRNs constrained by a non-zero viability and reproductivity implies that, for any 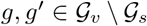, there exists a sequence of mutations which transforms *g*′ to *g* through GRNs in 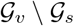. Since we presume that any single-locus mutation can occur with a non-zero probability (recall from section 2.1), there is a non-zero chance for *g*′ to mutate to *g* within one generation^2^, i.e., 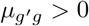. As a result, we observe that **T** is a positive matrix.

For the ease of presentation, we next show the convergence of equation (5) when the matrix **T** is symmetric and leave the proof for a non-symmetric **T** in Appendix D. In this case, the eigenvectors 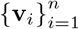 of the symmetric matrix **T** are linearly independent and form a basis of *n*-dimensional vectors, where 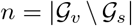. We order the eigenvectors such that the magnitudes of their corresponding eigenvalues 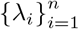 are non-increasing. The initial distribution can then be rewritten as a linear combination of the eigenvectors of **T**

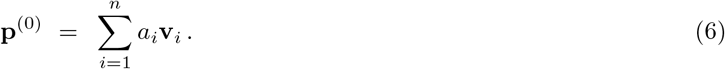

In addition, because **p**^(*t*)^ is proportional to **T p**^(*t*−1)^ for *t* > 0, we have **p**^(*t*−1)^ proportional to **T**^*t*−1^ **p**^(0)^ and consequently

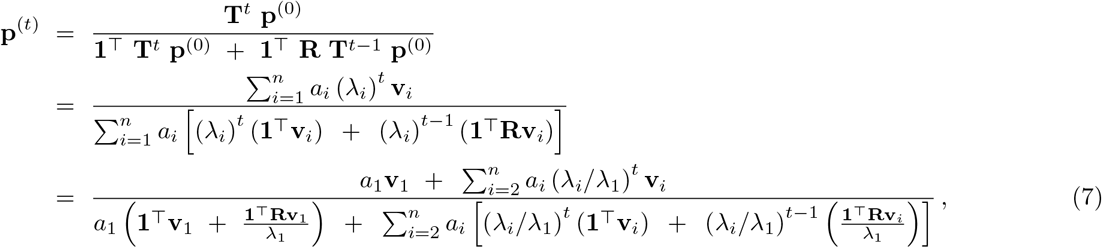

where **v**_1_ and *λ*_1_ are the leading eigenvector and the leading eigenvalue of **T** respectively. Since **T** is a positive matrix, by the Perron-Frobenius theorem, we have |λ_1_| > |λ_*i*_| for every *i* > 1, which guarantees the convergence of equation (5)

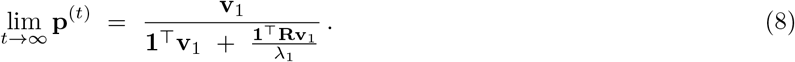

For a general and potentially non-symmetric matrix **T**, we can first factor **T** by its generalized eigenvectors and its Jordon normal form and then an analogous derivation follows (see Appendix D). Therefore, under the pathway framework of GRNs, the master equation (5) converges to a stationary distribution that is proportional to the leading eigenvector of **T**. Combined with the GRNs with a zero viability/reproductivity, whose probability to be observed under the limit *t* → ∞ can be easily computed given (8), the stationary distribution of GRNs describes the balanced scenario between selection, mutation, and reproduction.

### 3.2 Case Study: Binary Viability, Identical Reproductivity, and Independent Mutation

We next turn to a case study to validate our predicted stationary distribution of GRNs. We will examine a more specific version of the quasi-species model (4) with assumptions on the viability, reproduction, and mutation of a GRN. First, a GRN *g* either always survives the selection or becomes inviable, i.e., it has a binary viability *v*_*g*_ ϵ {0, 1}. It also implies that for any GRN 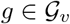 with a non-zero viability, we have *v*_*g*_ = 1.

Second, we assume that each GRN 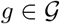 produces the same number of offspring and there is no sexual selection. Equivalently, the probability that an individual randomly sampled from an infinitely large offspring population is reproduced by a viable parent with GRN *g* is a constant for any viable GRN 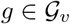. We denote this uniform reproductivity by *ρ*_*g*_ = ℙ[*I*_*t*−1_ → *I*_*t*_ | *g* (*I*_*t*−1_) = *g,* Ψ (*I*_*t*−1_)] = *ρ*, which is asserted to be non-zero.

Third, given the underlying collection of proteins Ω and genes Γ, the per-generation occurrence of mutation at every *γ* ϵ Γ is assumed an independent identically distributed Bernoulli random variable with a constant success probability *μ*. Moreover, if it occurs, a mutation at *γ* randomly changes *γ*’s expression behavior to any other pair of protein activator/product encoded in Ω with an equal probability. Under this assumption of independent and uniform mutation, the per-generation probability that a GRN *g*′ mutates to *g* becomes

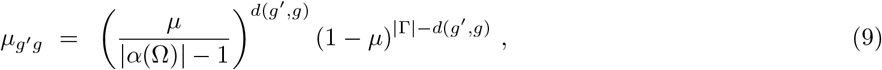

where we denote by *α*(Ω) the set of possible pairs of protein activator/product in Ω, and *d*(*g*′, *g*) is the number of genes with different expression behavior between *g*′ and *g*.

For this more specific model, we can rewrite the semi-transition matrix **T** into a series

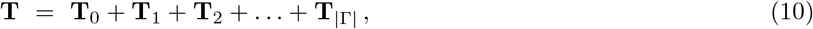

where the entry at the *i*_*g*_-th row and the 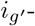th column of matrix **T**_*k*_ is 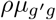 if *d*(*g*′,*g*) = *k* and 0 otherwise. Observe that **T**_0_ is proportional to the identity matrix **I** (of a proper size), and **T**_1_ is proportional to the adjacency matrix of the neutral network of GRNs (see section 2.3), which we denoted by **A**. Writing 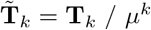, whose entries are finite even for a zero per-generation, per-locus mutation probability *μ*, equation (10) becomes

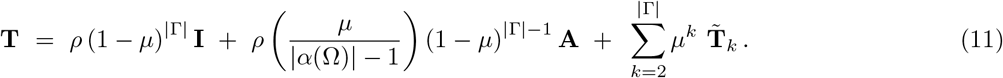

We further consider the scenario that mutations are rare events, specifically, under the limit *μ* → 0. Since the eigenvectors of **I**+*c***A** are exactly the eigenvectors of **A** for any scalar *c*, and 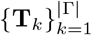 are symmetric matrices because *d*(*g*′,*g*) = *d*(*g,g′*), the theory of eigenvalue perturbation [54, 55] ensures that the leading eigenvector of **T** converges to the leading eigenvector^3^ of **A**:

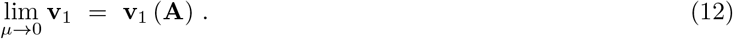

From equation (8), we have

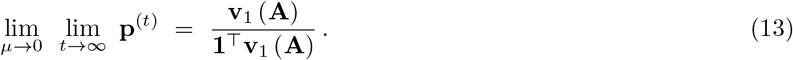

In network science, entries of the leading eigenvector of the adjacency matrix of a connected, undirected graph is known as the eigenvector centrality [56, 57] of the nodes. As a result, under the assumptions of binary viability, identical reproductivity, and rare, uniform mutation, *the probability distribution of viable GRNs converges to a stationary distribution that is proportional to their eigenvector centrality in the neutral network*.

To validate the predicted probability distribution of GRNs under mutation-selection balance, we simulate the evolution of 10^7^ parallel populations. The simulations are parametrized with the constant sets of |Γ| = 4 genes and |Ω| = 6 proteins. We further presume that two proteins can not be the product of any expression behavior, whose presence state can only be stimulated externally and hereafter they are referred to the *input* proteins. We also presume that two other proteins only have direct physiological effects and they can not serve as the activator of any expression behavior, which we call the *output* proteins. Under this minimal setup, there are in total |*α*(Ω)| = 16 potential pairs of expression activator/product, which leads to 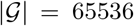 plausible GRNs. We evolve the populations under the environmental condition such that one of the input proteins is externally stimulated, and one of the output proteins shows a fatal effect which is required absent for an individual’s viability, resulting in 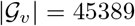 possible viable GRNs altogether. Our simulated GRNs are indeed small to avoid data scarcity in sampling the empirical distribution since the number of possible GRNs grows super-exponentially with the number of genes and proteins. We leave simulation of more realistic GRNs in future work.

The evolution of parallel populations are simulated using a Wright-Fisher model [58]. Specifically, we fix a number of 16 individuals for all populations, and given a current generation, the next generation is generated through randomly choosing viable GRNs from the current generation without replacement followed by potential mutations with a per-locus mutation probability *μ* = 0.1. We begin with 10,000 different initial populations where the GRN of every individual is chosen uniformly at random from all possibilities 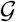, and 1,000 lineages are evolved from each initial population. Each of the 10^7^ parallel populations are evolved for a constant number of generations, from this ensemble of lineages we randomly sample a viable GRN to form the simulated distribution of GRNs. This fixed length of evolution is determined through the temporal lower bound such that the resulting GRN distribution is theoretically closed enough to the stationary distribution regarding to a given level of error tolerance (detailed in Appendix E).

Moreover, in order to account for the uncertainty of finite-sized sampling in the simulated distribution, we also draw the same number of 10^7^ independent samples from the predicted distribution (13) to form an empirical distribution. Repeating the sampling procedure 1,000 times, we obtain an ensemble of empirical distributions that captures the effect of finite-sized sampling over the predicted probability that GRNs are to be observed. We further use the averaged variation distance between the empirical distribution and the predicted distribution as the error tolerance from which the number of generations to be simulated is calculated such that convergence of the model is theoretically guaranteed (Appendix E).

In Figure 3a, we compare the exact, properly normalized leading vector of the transition matrix **T** (11) along with the predicted stationary distribution of viable GRNs under the rare-mutation approximation (13). Observe that even a moderate per-locus mutation probability *μ* leads to a GRN distribution well aligned with the predicted one, especially, with respect to the uncertainty arising from finite-sized sampling in the simulations. Moreover, Figure 3b shows the simulated distribution of viable GRNs after long-term evolution. We see that, despite a little overdispersion, the simulated distribution agrees with the derived stationary distribution of GRNs. Direct comparison between the simulated distribution and the exact solution, i.e., the leading eigenvector of the transition matrix (11) shows no significant difference as well (see Supplementary Figure 5). Combined, our simulations indicate computational evidence that, when viability is assumed rugged and mutations are rare, the topology of the neutral network, particularly the eigenvector centrality of mega-nodes, serves as a informative predictor of the prevalence of GRNs under mutation-selection balance.

**Figure 3:**
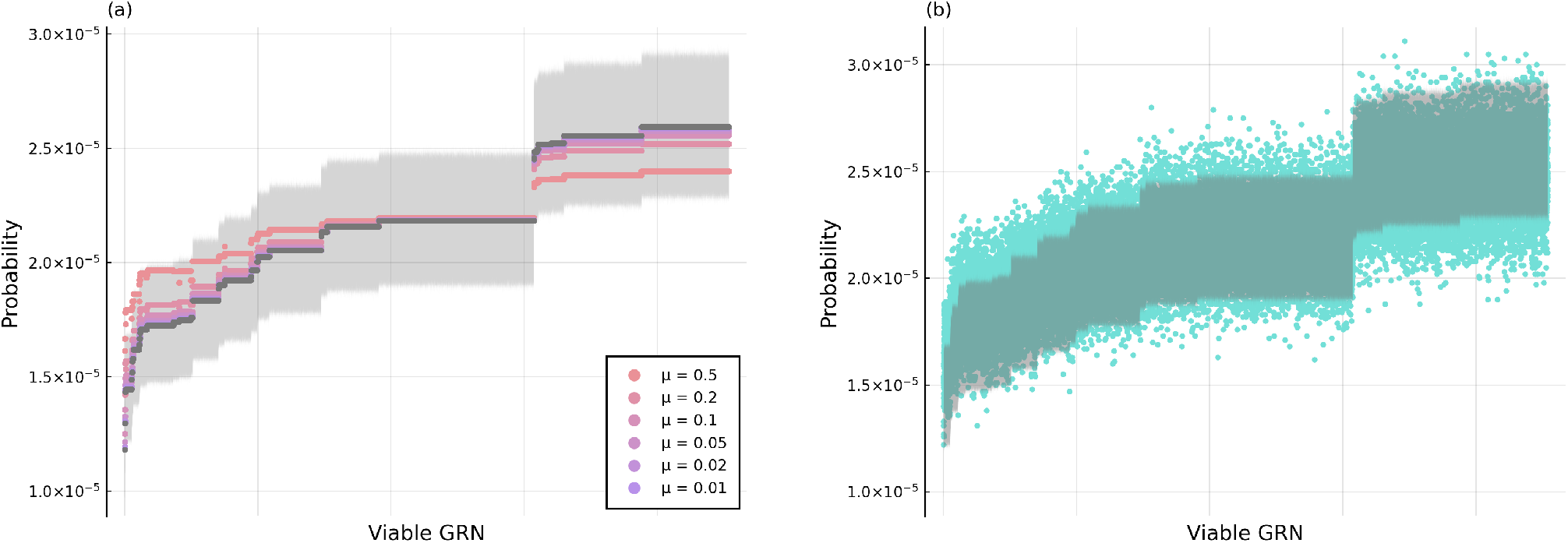
Validation that the evolutionary dynamics of GRNs converges to the derived stationary distribution (13). We compare the predicted stationary distribution of viable GRNs under the rare-mutation approximation with (a) the exact leading eigenvector of the transition matrix (11) with various per-locus mutation probability *μ* (colored by a red-purple gradient from large to small), and (b) the distribution of GRNs sampled from their simulated evolutionary dynamics with *μ* = 0.1 (blue). The predicted distribution is colored in gray, and the shaded area shows its 95% confidence band that accounts for the uncertainty of finite-sized sampling in the simulations. In both panels, the viable GRNs are ordered increasingly by their predicted probability to be observed.

### 3.3 Prevalent GRNs under Mutation-Selection Balance

We now apply our prediction in the case study of binary viability, identical reproductivity, and rare mutation to further investigate the structure of GRNs that have a higher probability to be observed than others under different environmental conditions. Here we again consider GRNs with a constant collection of 6 proteins and 4 genes. In addition, for the ease of presentation, we label the genes by uppercase letter Γ = {*A, B, C, D*} and the proteins by numerals Ω = {1, 2, 3, 4, 5, 6}, where protein 1 and 2 are the input proteins and protein 5 and 6 are the output proteins respectively (see section 3.2). Under the pathway framework of GRNs, an environment can be jointly described by a.) a set of stimuli proteins that are externally stimulated to be in the presence state, b.) a set of essential proteins whose absence state leads to inviability of the individual, and c.) a set of fatal proteins whose presence state also causes inviability. We will focus on seven distinct environments listed in Table 1 that showcase the scenarios of single versus multiple stimulated/essential/fatal proteins and their combinations.

**Table 1:**
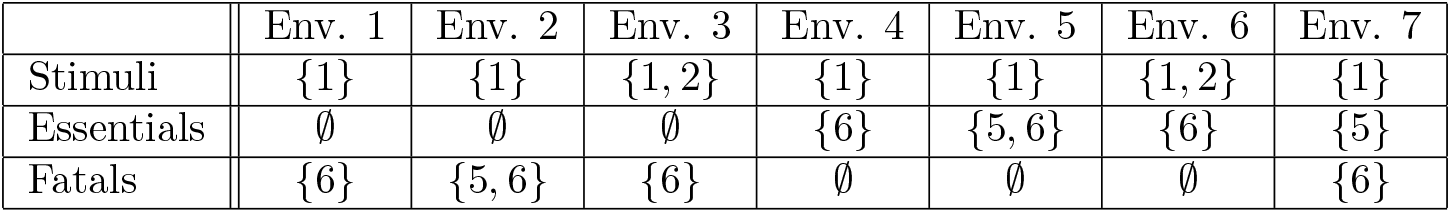
Different environmental conditions specified by the sets of stimulated, essential, fatal proteins.

For each of the focal environmental conditions, we examine the prevalent regulatory structure among various groups of GRNs. These groups consist of GRNs satisfying different constraints on their structural properties, which correspond to a few artificially enforced scenarios of interests. Here the focal topological constraints originate from patterns observed in the most prevalent GRNs, and progressively adding constraints offers a rough ranking of regulatory patterns for their inducing prevalence. We arrange groups of GRNs based on the following four constraints: First, GRNs with a gene of “spare” functionality are excluded, where the spareness of a gene refers to its negligible consequence on the resulting phenotype. This includes self-regulating genes due to the binary state assumption and genes that are activated by an input protein which is not externally stimulated or that produce an output protein without an essential/fatal effect under the given environment. Second, we exclude GRNs with multiple genes of the same, redundant expression behavior. Third, we only consider those GRNs where all the genes are functionally activated. This constraint mimics the scenario that genes with active expression behavior are more likely to be observed empirically than inactive ones. Forth, we exclude GRNs where a gene is directly activated by a stimulus and produces an essential protein to enforce selection on regulation rather than individual genes. Combinations of these four constraints lead to eight distinct groups where the prevalent GRNs are investigated (see Table 2).

**Table 2:** Groups of GRNs by imposing constraints on their structural properties.

|  | No spare genes | No redundant genes | All genes activated | No direct selection |
| --- | --- | --- | --- | --- |
| Group (i) |  |  |  |  |
| Group (ii) | ✓ |  |  |  |
| Group (iii) | ✓ | ✓ |  |  |
| Group (iv) | ✓ |  | ✓ |  |
| Group (v) | ✓ | ✓ | ✓ |  |
| Group (vi) |  | ✓ |  |  |
| Group (vii) |  |  |  | ✓ |
| Group (viii) |  | ✓ |  | ✓ |

In Figure 4, we plot the GRNs that have the largest predicted probability to be observed among the various groups and environments, i.e., the GRNs with the greatest eigenvector centrality in the neutral network under each scenario. Note that such GRNs may not be unique; in fact, one can find multiple alike GRNs through transformations that preserve their roles in the neutral network, e.g., exchanging the expression behavior of two genes *A* and *B*. Yet, these GRNs all share the common structural features, and we only show a random sample from the GRNs with the same, maximal probability to be observed in our prediction. Moreover, Figure 4 demonstrates the prevalent GRNs in both the representation of the pathway framework that manifests expression activator/product of each gene (labeled arrows between circles) and that of the conventional notion showing the regulation between genes (unlabeled arrows among rectangles).

**Figure 4:**
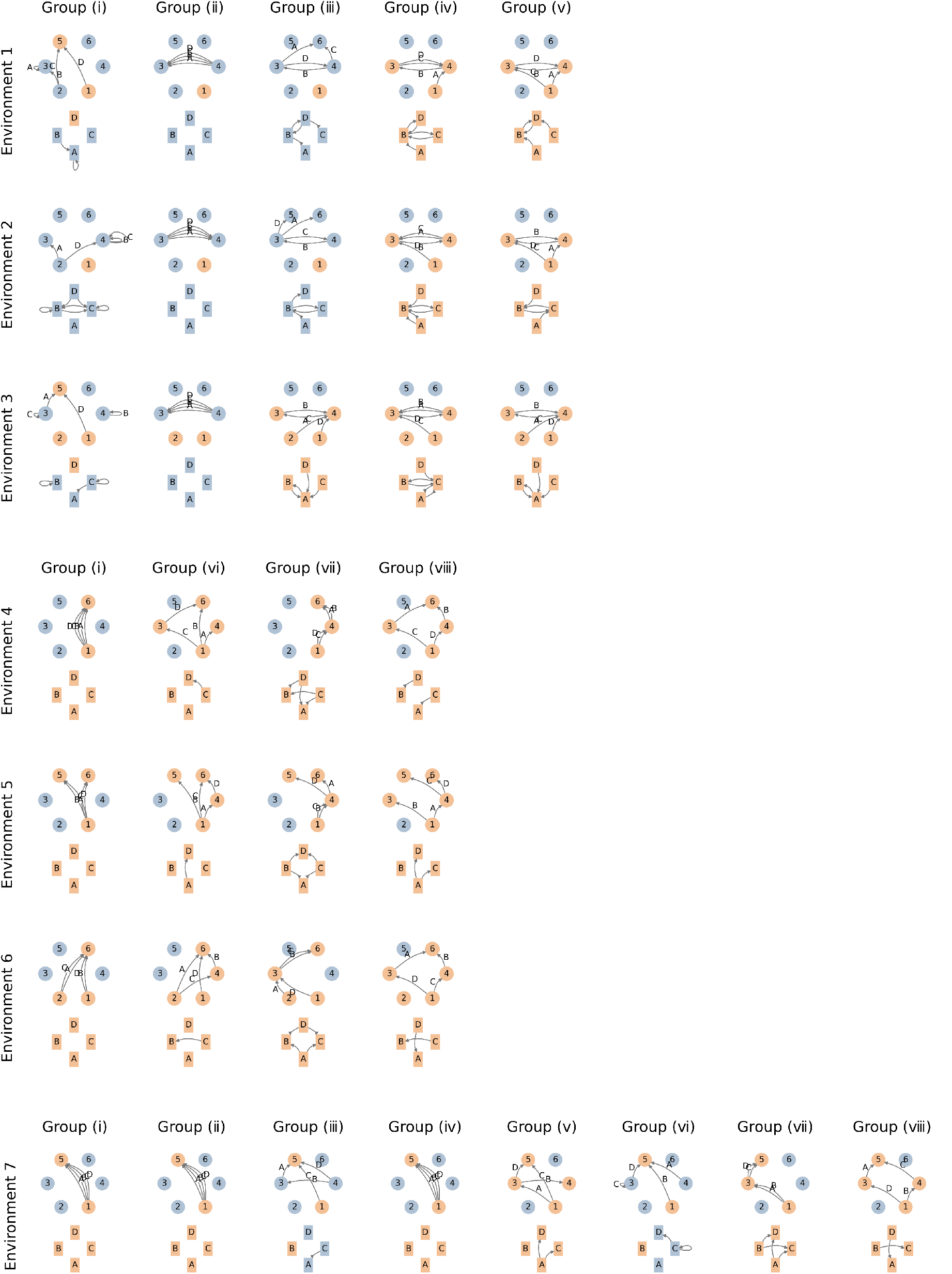
GRN that has the largest eigenvector centrality in the neutral network for different environmental conditions (Table 1) and among different constrained groups of GRNs (Table 2). For each prevalent GRN, its pathway framework representation is plotted by the circles and the labeled arrows, while its conventional representation is drawn through the rectangles and the unlabeled arrows. A node is colored in orange if the protein/gene is present/activated and in blue otherwise.

A few intriguing observations arise from the prevalent regulatory structure in Figure 4. For the environmental conditions where only the fatality of protein products is imposed (environment 1, 2, and 3), the GRNs with the largest probability to be realized under mutation-selection balance are the ones in which spare genes dominate (group (i)). Once constrained by the absence of the spare genetic functionality (group (ii)), we see prevalent GRNs favoring all genes sharing the same expression behavior which does not involve any stimulated or fatal proteins. If we further exclude redundant genes or enforce all genes to be activated (group (iii) and (iv) respectively), the prevalent GRNs demonstrate a structure which seemingly avoids expression activated by the stimulated protein or producing the fatal proteins as much as possible, and imposing both constraints leads to a similar outcome. Interestingly, for the environment with multiple stimuli and constraining on no spare and redundant genes (environment 3 and group (iii)), the functional activeness of all genes naturally emerges.

On the other hand, for the environmental conditions where only the essentiality of protein products is obligated (environment 4, 5, and 6), the most prevalent GRNs are the ones where several redundant genes are directly activated by a stimulus and produces an essential protein, and they are evenly split if multiple essential targets or stimuli exist (group (i)). When redundant genes are artificially excluded (group (vi)), the prevalent GRNs turn into a structure that manifests multiple pathways between the stimuli and the essential proteins. While constrained by no direct gene expression activated by a stimulus and producing an essential target (group (vii)), the prevalent GRNs similarly show multiple pathways yet each of which involves at least two genes, and these pathways share the same intermediate protein that serves as the product of one and the activator of another. Jointly imposing the two constraints mentioned above (group (viii)) results in the prevalent GRN structure that maintains multiple regulatory pathways and simultaneously triggers the presence state of the underlying proteins, if plausible. Notice that for these environmental condition, the prevalent GRNs take advantages of the functionality of every gene, and all the genes are activated.

Last but not least, for the environmental condition where both essential and fatal proteins exists (environment 7), the most probable GRNs favor redundant genes that are directly activated by the stimulus and synthesize the essential target when the genetic redundancy is not constrained (group (i), (ii), and(iv)). Otherwise, the prevalent regulatory structure leaves one gene to maintain its essentiality, whereas others are capable to generate the essential protein but their activators remain absent (group (iii) and (vi)). If we further artificially require the activation of genes or exclude direct selection on individual genes (group (v), (vii), and (viii)), we begin to see multiple pathways in the prevalent GRNs.

We discover that most of the prevalent structures of GRNs in Figure 4 follow an intuitive pattern: these GRNs have the least plausible, subsequent inviable mutants under the diverse environmental conditions and structural constraints. When selection is enforced by the fatality of proteins, any regulatory pathway from the stimulus to the fatal protein is prohibited. The prevalent GRNs keep the fewest proteins in their presence state that can serve as the potential expression activators, since this minimizes the ways for subsequent mutations to create a lethal regulatory pathway. As a result, the mutation-selection balance drives the dominance of genes with a spare functionality, and secondly the appearance of redundant genes whose expression involves neither the stimulus nor the fatal protein. On the contrary, when selection acts through the essentiality of proteins, regulatory pathways from the stimulus to the essential target become critical for an individual’s viability. The prevalent GRNs show the structure of redundant genes or multiple pathways such that the chance of eliminating the essential pathways through subsequent mutations is most mitigated. Therefore genes in these prevalent GRNs are expected to be functionally active. Moreover, even if a gene does not participate in an essential pathway, its expression behavior will involve the stimulus or the essential protein to potentially form a pathway with latent mutations. In the case where both the fatal and the essential target exist, the prevalent GRNs demonstrate structures as a superposition of the two patterns we previously discussed, which alternatively display the characteristics of the fatality-/essentiality-driven scenario under different structural constraints of gene regulation.

## 4 Discussion

In this work, we analyze the evolutionary dynamics of GRNs under a quasi-species model with selection, mutation, and asexual reproduction. Integrating with the pathway framework of GRNs that abstracts the alleles of genes through their expression behavior, we analytically show that the population dynamics always converges to a stationary distribution of GRNs given any arbitrary viability function and stochastic mutational transition as long as no mutation is prohibited. This stationary distribution characterizes the ensemble of regulatory circuits under mutation-selection balance, and it implicates the structural features of GRNs to be predicted favorable under long-term evolution. Next, we investigate a case study assuming binary viability, identical reproductivity and rare mutation, and find that the stationary distribution of GRNs can be derived from the topology of the genotype network. Specifically, the probability to observe a GRN under mutation-selection balance is proportional to the GRN’s eigenvector centrality in the neutral network, which is a subgraph of the genotype network consisting of all viable GRNs.

We advocate that our contribution to the theory of evolutionary dynamics provides a mechanistic explanation for the key assumption of irreducible transition matrices in existing literature [45, 46, 47]. As we mentioned in the Introduction, Moran relates this assumption, which leads to the global convergence to a single quasi-species, to the scenario that viable genotypes are mutually accessible through mutations [48]. In a similar spirit, a recent review concludes that a population genetic model situating on a genotype network always evolves into a stationary solution once we retreat to the regime of non-zero fitnesses. Our work takes an alternative route to recover the same assumption; instead of the absent knowledge between genotypes and their fitnesses, we consider a minimal modeling framework of how genotypes develop into phenotypes via the mechanisms of gene regulation [37]. Despite the simplicity of this modeling framework, when encapsulating the genotype-phenotype mapping through GRNs, the mutational accessibility between genotypes with non-zero fitnesses naturally emerges due to the high dimentionality of GRNs. This result relaxes the global convergence in quasi-species theory to the cases with extreme fitness values such as a holey adaptive landscape [59].

Our derivation sheds light on how we may interpret the prevalence of GRNs under rare mutation and strong selection on the resulting phenotypic functionality. When first introduced [56], the eigenvector centrality was designed to capture an individual’s global “importance” as measured by their social ties in a communication network. In particular, the eigenvector centrality is computed based on the idea that a node’s importance is proportional to the sum of its neighbors’ importance scores. This interpretation is nicely translated to the content of the neutral network of regulatory circuits: Under mutation-selection balance, our derivation predicts that the probability to observe a GRN is proportional to the total likelihood to find its viable, mutational neighbors in the population. Intriguingly, the interpretation of eigenvector centrality leads to some emerging concept of robustness [60], where the prevalence of a GRN is not only due to its selective advantage but also the overall prevalence of its mutational neighboring GRNs.

Moreover, the observed prevalent structures of GRNs in our analyses also provides a possible alternative explanation for evolutionary robustness. We inductively find that these prevalent regulatory structures follow the same pattern to achieve a minimal number of plausible inviable mutants (see section 3.3). Since the genotype network is a regular graph under the pathway framework of GRNs (recalling from section 2.3), i.e., every GRN has the same amount of mutational neighbors, minimizing the number of inviable mutants optimally increases the viable mutants for a GRN. In other words, the observed prevalent GRNs under various environmental conditions appear to show the regulatory structures with the maximal number of neighbors in the neutral network, and indeed the degree of a node is known to be strongly correlated with its eigenvector centrality in the network science literature [57]. We emphasize that these concepts of robustness naturally emerge from the mechanistic, quasi-species model of GRN evolution rather than an *a prior* assumption about prevalent regulatory circuits.

Previous work often focused on relating the topological features of genotype networks to evolutionary processes of interest. For example, evolvability has been approximated by the size of the genotype network of a given phenotype [61], as well as the number of “neighboring” phenotypes inferred from the genotype network [62]. Robustness has been modeled as the node degree in the genotype network [62], and [63] adopted the average path length in the genotype network as a proxy for genetic heterogeneity. To our best knowledge, Van Nimwegen et al. was the first to bridge between the asymptotic abundance of different genotypes under a population genetic model and their eigenvector centrality in the neutral network [60]. Our case study resonates Van Nimwegen et al.’s conclusion and differentiates a quasi-species perspective from a model lacking of genetic variation in a population. For instance, if a population fixes a single genotype at all time and its evolution is modeled as a random walk on the neutral network, network science guarantees the fixation probability at a given genotype to be proportional to its *degree* instead of the eigenvector centrality in the neutral network [57].

The current scope of the work presented here is not without a few noteworthy limitations. First, we assume a constant, static surrounding in which the population evolves, whereas populations certainly experience shifting or alternating environmental conditions [64, 65, 66]. Second, our model mainly focused on the joint forces of selection and mutation. Although this simple model can indeed be extended through more sophisticated mechanisms known to play a role in evolutionary dynamics such as recombination [67, 68], gene duplication [69, 70], and demographic information [71], we leave such extensions–along with their possible implications–to future work. Third, when the time scale of environmental changes is much faster than that of the evolutionary dynamics (see Appendix E), the transient constitution of GRNs in a population shall acquire more attention than their stationary distribution at mutation-selection balance [72, 73, 74]. Put simply, it remains an open question whether real-world populations should ever be conceptualized as at equilibrium (even dyanmic) as opposed to existing in some far-from equilibrium state [75]. As a result, further investigation should focus on the transient distributions and/or trajectories of GRNs under various population genetic models. Finally, despite confirmation between the derived stationary distribution of GRNs in an infinitely large population and the long-term numerical simulations, we also find that a finite population size moderately influences the transient evolutionary dynamics. Developing a richer understanding of the role drift plays in structuring the evolution of GRNs is an important extension of our work.

The observed structure of molecular interaction networks is a result of myriad evolutionary forces. By analyzing such topologies using a network-science approach, it may be possible to construct a mechanistic theory for how evolution shapes and is constrained by higher-order interactions. Across a broad scope of genotype/neutral networks– with applications ranging from RNA sequences to metabolic reactions–our work rigorously shows that the neutral network of GRNs must be connected (in agreement with existing computational work [76]) and that the relative frequency at equilibrium of various GRNs can be predicted from first principles. Therefore, our work connects the evolutionary forces/mechanisms embedded in a population genetic model with the accordingly favorable GRN structure through the topology of the neutral network. Clearly, our predicted prevalent regulatory structures under mutation-selection balance may not capture all the features in empirical GRNs [77, 78]; however, we establish a null expectation for how GRNs are shaped by mutations and selection [26, 27]. Critically, this null expectation appears to recapitulate many of the topological features of molecular interaction networks currently associated with evolvability and robustness. Perhaps, more broadly speaking, the emergence of complex fitness landscapes can result from simple evolutionary rules.

## Acknowledgement

Hold.

## Data Availability

The authors affirm that all data necessary for confirming the conclusions of the article are present within the article, figures, and tables.

## Appendices A Mathematical Formulation of the Pathway Framework of GRNs

In this study, we assume there is a constant set of proteins that can *possibly* appear in the organisms. We clarify that this constant collection is not necessarily the proteins which we have observed in the certain species to date; contrarily, these proteins are the plausible options of the activators and products of gene expression, and they are better acknowledged as all (or a reasonable subset of) the proteins under our awareness. We will refer to the *state* over this protein set for their actual appearance in the organisms, with a more detailed discussion later.

We furthermore divide the constant set of proteins into three categories: *input proteins* that can only be supplied through external stimuli but not through any internal gene expression, *output proteins* that are products of gene expression which affect physiological traits of the organisms but can not serve as activators of gene expression, and the remaining *internal proteins* with neither constraints. The input and internal proteins form the set of plausible activators for gene expression, whereas the internal and the output proteins become the set of products. This completes the underlying backbone of GRNs under the pathway framework.

### Definition A.1.

We denote by Ω_*s*_ and Ω_*t*_ be the fixed underlying **activator set** and **product set** of gene expression respectively. And we call their union Ω = Ω_*s*_ ∪ Ω_*t*_ the underlying **protein set**.

The three categories of proteins can be recovered easily from the notion of expression activators and products. In particular, the input, output, an d internal proteins are Ω − Ω_*t*_, Ω − Ω_*s*_, and Ω_*s*_ ∩ Ω_*t*_ respectively.

With a pre-specified underlying backbone (Ω_*s*_, Ω_*t*_) of the regulatory structures, a gene regulatory network is a graphical abstraction of the expression behavior for the whole genotype. We will assume that the collection of genes of the organisms remains the same over evolutionary time, i.e., there is no duplication and deletion of the loci. A GRN is then uniquely determined by the activator and the product of every gene, and we have the following formulation:

### Definition A.2.

Denote by Γ the fixed set of genes, or the **gene set**. We define a **gene regulatory network (GRN)** as a mapping *g* : Γ → Ω_*s*_ × Ω_*t*_. We further denote by 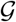 the set of all such gene regulatory networks.

Definition A.2. may seem an unusual way to describe a network. To illustrate that the definition is appropriate, recall that a directed edge in a GRN under the pathway framework represents the input-output relation of a gene’s expression. Every edge in the GRN is thus labeled by the gene whose allelic content is abstracted as the edge. With the given protein set Ω as nodes, the GRN can be described as its edgelist representation — a table where each row stands for an edge (and its corresponding gene) and the two columns entails its source and target (i.e., the corresponding activator and protein product respectively). This table, and therefore the GRN, is equivalent to a mapping *g* from the finite set of genes Γ to the pairs of activators and products Ω_*s*_ × Ω_*t*_, where *g*(*γ*) is the expression input-output pair of gene *γ* ϵ Γ.

We also have the notion of projection from the input-output relations of a genotype. For any gene *γ* ϵ Γ, these projections explicitly point to its activator protein *s*_*g*_(*γ*) and its protein product *t*_*g*_(*γ*):

### Definition A.3.

The **activator projection** and the **product projection** of a gene regulatory network *g* are defined by *s*_*g*_ = *p*_*s*_ ○ *g* and *t*_*g*_ = *p*_*t*_ ○ *g*, where *p*_*s*_ and *p*_*t*_ are the set projection from Ω_*s*_ × Ω_*t*_ onto Ω_*s*_ and Ω_*t*_ respectively.

We next introduce how the two evolutionary forces we will consider in a population genetic model of GRNs, mutation and selection, can fit into the pathway framework.

Mutating the allele of a gene can alter the expression behavior of the gene. Since under the pathway framework a genotype is conceptualized as a GRN on the expression functional level, we will model mutation to be changing the input-output relation of gene expression. Specifically, a mutation randomly rewires a single edge in the GRN and results in a mutant GRN. Equivalently we can find all the possible mutants, namely, those only differ by one input-output pair from the original GRN, and a mutation can be defined as a random process over the mutants.

### Definition A.4.

Let *g*_1_, *g*_2_ be two gene regulatory networks. The set of genes with different alleles between *g*_1_ and *g*_2_ is

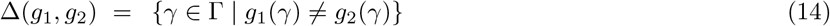

The **edit distance** between *g*_1_ and *g*_2_ is defined by *d*(*g*_1_, *g*_2_) = |Δ(*g*_1_, *g*_2_)|.

### Definition A.5.

Let *g* be a gene regulatory network. We denote the set of **mutants** from *g* by

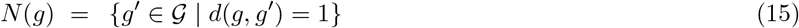

i.e., those gene regulatory networks that are 1-edit-distant from *g*.

### Definition A.6.

A **mutation** of gene regulatory network *g* is a random process with a probability measure over its mutants *N* (*g*), which we denote by *μ*_*g*_.

On the other hand, natural selection can be regarded as a phenotypic response to the surrounding environment, where the phenotype is derived from the genotype. We presume that the physiological traits of an organism are uniquely determined by the actual appearance of proteins within it, and that they are conditionally independent of the external environments. The phenotype is thus the collective state over the underlying protein set Ω. And this collective state is the outcome of external environmental stimuli and internal chemical signals propagating on the gene regulatory networks.

For simplicity we adapt the chemical state of proteins to be binary, i.e., that a protein is present in the organism versus that it is absent. Additionally, assuming that the environmental condition directly triggers the presence state of some proteins, the binary state of a protein is determined by its reachability from those stimulated ones on the GRN. We let the set of proteins with the presence state to represent the phenotype derived from the GRN:

### Definition A.7.

Let a *g* be a gene regulatory network, and let Ω_0_ ⊂ Ω−Ω_*t*_ be the set of environmentally stimulated proteins. The **phenotype** of *g* is the function 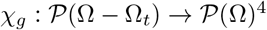, where for any protein *ω* ϵ Ω, *ω* ϵ *χ*_*g*_(Ω_0_) if and only if there exists a sequence of genes 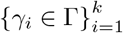 such that *t*_*g*_(*γ*_*k*_) = *ω*, *s*_*g*_(*γ*_*i*+1_) = *t*_*g*_(*γ*_*i*_) for *i* = 1,2, … , *k* − 1, and *s_g_*(*γ*_1_) ϵ Ω_0_.

The phenotypic response to the environmental condition, or namely the individual viability under natural selection, becomes a function of the collective binary state of the underlying proteins Ω. We again for simplicity adapt the viability to be the binary variable that whether the individual organism survives or not. Moreover, we suppose that this binary viability solely depends on two collections of proteins: those proteins which are essential for the organism to survive, and those having fatal effects to the organism. The selective environment is then explicitly specified by the sets of stimulated, essential, and fatal proteins respectively. We describe the outcome of selection through the viable GRNs, i.e., those with which a organism will survive natural selection:

### Definition A.8.

Let Ω_0_ ⊂ Ω − Ω_*t*_ and Ω_+_, Ω_−_ ⊂ Ω − Ω_*s*_ be the stimulated, essential, and fatal proteins in the environmental condition respectively. The selective environment, or simply **selection**, is the triplet 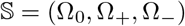. We define the set of **viable** gene regulatory networks under selection 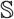 by

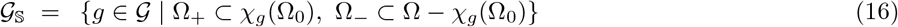

Notice that we have implicitly exerted the assumption that the stimulated proteins must be a subset of the input proteins, and the essential and fatal proteins must be a subset of the output proteins (recall Definition A.1).

## B Formal Definition of the Genotype Network and the Neutral Network of GRNs

With the constant sets of activators Ω_*s*_, products Ω_*t*_, and genes Γ, and the pre-determined sets of stimulated proteins Ω_0_, essential proteins Ω_+_, and fatal proteins Ω_−_, we have the following definitions:

### Definition B.1.

Recall from Definition A.2 and A.5 that 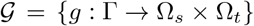 is the set of all plausible gene regulatory networks, and *N* (*g*) are mutants from gene regulatory network *g*. The **genotype network** is a graph *G* whose nodes 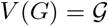 and whose edges 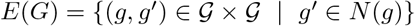.

### Definition B.2.

Let 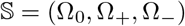 be a given selection, and recall from Definition A.8 that 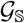 is the set of viable gene regulatory networks under 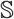. The **neutral network** subjected to 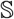 is a graph 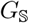 whose nodes 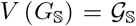 and whose edges 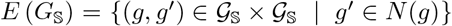. Note that 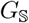 is the induced subgraph of the genotype network *G* on nodes 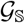.

## C Structural Properties of the Genotype Network and the Neutral Network of GRNs

We begin with analyzing the structural properties of the genotype/neutral network of GRNs under the pathway framework, as well as highlighting those that are relavent to deriving the stationary distribution in the generalized population genetic model. First of all, the genotype network *G* shows an intuitive and nicely ordered structure. Since the mega-nodes in *G* consist of all the plausible GRNs 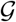 given the constant activators Ω_*s*_, products Ω_*t*_, and genes Γ, every mega-node is equivalent to a tuple of |Γ| entries, each of which takes a discrete value from Ω_*s*_ × Ω_*t*_. Two meganodes/GRNs are connected in *G* if and only if they differ by the allele of a single gene, namely that the two corresponding tuple only differ by one entry, and as a result, the genotype network *G* is essentially a high-dimensional lattice.

The lattice-like nature of the genotype network *G* implies several structural properties. The genotype network *G* must be a connected graph, which agrees with the intuition that any two genotypes (at least on their gene expression level, i.e., the GRNs) can be mutually reached by a sequence of mutations under zero selection pressure. In addition, the distance between two GRNs *g*_1_ and *g*_2_ in *G* is, recalling from Definition A.4, exactly their edit distance *d* (*g*_1_, *g*_2_) because the shortest paths correspond to the scenarios to mutate the genes with different alleles Δ (*g*_1_, *g*_2_) sequentially. Furthermore, we also see that any GRN has the same number of mutational neighbors in *G*:

### Lemma C.1. The genotype network G is a regular graph.

*Proof.* Given an arbitrary gene regulatory network 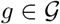 and for any gene *γ* ϵ Γ, there are |Ω_*s*_ × Ω_*t*_| − 1 other gene regulatory networks that only differ from *g* by the allele at *γ*. The number of mutatnts is

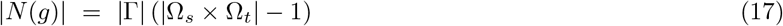

for any gene regulatory network 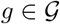, and hence every mega-node in *G* has the same degree.

On the other hand, although the neutral network 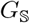 subjected to a pre-determined selection 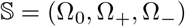 is a subgraph of the genotype network *G* (see Definition A.8), its structure is more disordered. There is no guarantee that 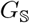 is regular, and in fact one can easily find some counter-examples (e.g., see Figure). The distance between two GRNs in *G* may not be preserved in 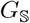 either. For example, consider the case where Ω_*s*_ = {1, 2, 3}, Ω_*t*_ = {2, 3, 4}, Γ = {*a, b*}, Ω_0_ = {1}, Ω_+_ = 4 and Ω_−_ = ∅, and two GRNs *g*_1_ and *g*_2_ such that *g*_1_(*a*) = (1, 2), *g*_1_(*b*) = (2, 4), *g*_2_(*a*) = (1, 3) and *g*_2_(*b*) = (3, 4). In the genotype network *G*, there are two length-2 paths between *g*_1_ and *g*_2_, either through GRN *g*_3_ or *g*_4_ where *g*_3_(*a*) = *g*_2_(*a*), *g*_3_(*b*) = *g*_1_(*b*), *g*_4_(*a*) = *g*_1_(*a*) and *g*_4_(*b*) = *g*_2_(*b*). However, neither *g*_3_ nor *g*_4_ satisfy the selection criterion, and thus they are excluded from the neutral network 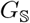, in which the distance between *g*_1_ and *g*_2_ is greater than 2.

Nevertheless, it turns out that, in most scenarios, any two GRNs are mutually reachable through some mutational trajectory in the neutral network 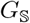:

### Lemma C.2.

If |Γ| > |Ω_+_|, then the neutral network 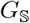 under selection 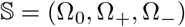 is a connected graph.

*Proof.* To show that 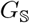 is a connected graph, our strategy follows: For any two viable gene regulatory networks 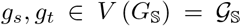, we will find a sequence of viable GRNs 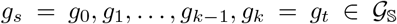 that form a mutational trajectory from *g*_*s*_ to *g*_*t*_, i.e., 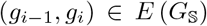 for every *i* = 1, 2, *… , k*. More specifically, we will uncover the sequence of mutations through a few general “steps” of edge rewiring in GRNs and ensure that two invariants hold in each of these steps:

I. There is a path from the stimulated proteins Ω_0_ to each of the essential proteins Ω_+_ in the GRNs.
II. There is no path from the stimulated proteins Ω_0_ to any of the fatal proteins Ω_−_ in the GRNs.

Here we would like to introduce a few notations for the ease to illustrate the edge-rewiring steps in the GRNs. First, we put the genes into different groups with respect to *g*_*s*_ and *g*_*t*_. Let 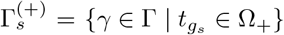 and 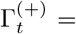 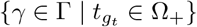 be the genes that directly produce the essential proteins in *g*_*s*_ and *g*_*t*_ respectively. Similarly, let 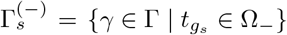 and 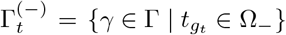 be the genes that directly produce the fatal proteins in *g_s_* and *g_t_* respectively. Graphically, these genes correspond to the incoming incident edges of either Ω_+_ or Ω_−_. In addition, denote by Π_*g*_(*u, v*) the set of genes that are involved in paths from protein *u* to protein *v* in the GRN *g*, and let 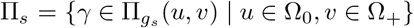 and 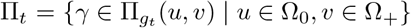 be the genes that are involved in pathways from Ω_0_ to Ω_+_ in *g*_*s*_ and *g*_*t*_ respectively.

Second, when Γ \ Π_*s*_ or Γ \ Π_*t*_ is non-empty, there exists some “safe” allele among all the plausible allelic contents Ω_*s*_ × Ω_*t*_. We will denote such a tuple as *α*. If Ω_*s*_ ∩ Ω_*t*_ is non-empty, then there is a protein *ω*^t^ that can serve both an activator and a product, and we will take *α* = (*ω*′, *ω*′). On the other hand, if Ω_*s*_ ∩ Ω_*t*_ = ∅, we must have a non-empty Ω_*s*_ \ Ω_0_, otherwise Γ \ Π_*s*_ = Γ \ Π_*t*_ = ∅. Hence there is a protein *u*′ ϵ Ω_*s*_ \ Ω_0_ of the absence state, and we will take *α* = (*u*′, *v*′) for some *v*′ ϵ Ω_−_. Note that the allele *α* is said “safe” in the sense that introducing *α* will never break invariant (II).

Now we state in details the five steps of edge rewiring that mutate *g*_*s*_ into *g*_*t*_ through viable GRNs:

1. Rewire edges (alleles of genes) in *g*_*s*_ to generate a viable GRN 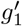 such that 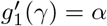 for any gene 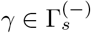 and 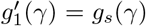 for 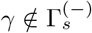. During this rewiring process, invariant (I) holds since the alleles of 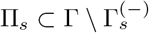 remain unchanged, and invariant (II) holds because this step simply introduces the safe allele *α*.
2. Rewire edges in 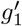 to generate another viable GRN 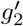 such that 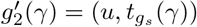 for any gene 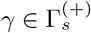 and some *u* ϵ Ω_0_, and 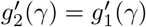 for 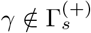. Since this step only creates length-1 pathways from Ω_0_ and Ω_+_, both invariant (I) and (II) are guaranteed.
3. Rewire edges in 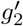 to generate another viable GRN 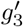 such that 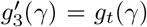 for any gene 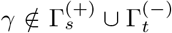 and 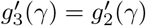 for 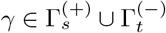. Invariant (I) holds because the length-1 pathways introduced in Step 2 remain unchanged. Since in 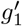 (and thus 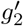) proteins Ω_−_ have no incoming incident edges, and no rewiring leads to an incoming edge of Ω_−_ in this step, invariant (II) is also ensured.
4. Rewire edges in 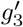 to generate another viable GRN 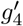 such that 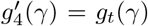 for any gene 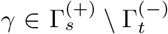 and 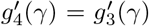 for 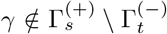. In particular, for a gene 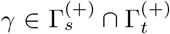, the rewiring process can be achieved via an intermediate gene *γ*′ ϵ Γ due to the pre-condition that |Γ| > |Ω_+_|^5^. Since for each 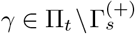, Step 3 has properly rewire its edge/allele to 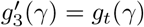, the essential pathways formed by Π_*t*_ are gradually completed throughout the process of rewiring edges corresponding to 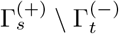. And similar to Step 3, no edge is rewired to be an incoming edge of Ω_−_ in this step, and thus invariant (II) holds as well.
5. Rewiring edges corresponding to 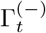 in 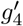 to generate *g*_*t*_ completes the viable mutational trajectory. Because edges of 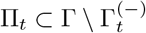 remain unchanged in this step, invariant (I) still holds. Furthermore, since *g*_*t*_ is viable, rewiring edges of 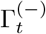 also preserves invariant (II).

Note that since for a GRN to satisfy the selection criterion, every protein in Ω_+_ must be produced by a gene, so we must have |Γ| ≥ |Ω_+_|. As a result, the only case that Lemma C.2 has excluded is that of |Γ| = |Ω_+_|, where the GRNs form |Ω_+_| ! components in the neutral network, each of size |Ω_0_|^|Ω_+_|^. Moreover, the proof we provide here is general enough such that Lemma C.2 holds even if one adapts additional constraints and defines a mutation as changing either the protein activator or the protein product of a gene but not both.

## D Convergence to a Stationary Distribution with a Non-symmetric Transition Matrix

Here we show a general, analogous proof for (8) in the case that the semi-transition matrix **T** is non-symmetric. Any square matrix can be factored by its general eigenvectors and its Jordan normal form. In particular, we have **T** = **PJP**^−1^ (or equivalently **TP** = **PJ**), where **P** is a matrix consisting of linearly independent column vectors, and **J** is a block diagonal matrix such that

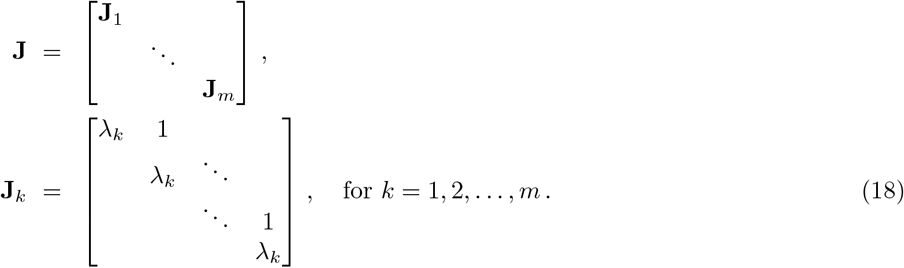

The diagonal entries of **J** are the eigenvalues of **T** with multiplicities. The matrix **J** is called the **Jordan normal form** of **T**, and the column vectors of **P** are called the **generalized eigenvectors** of **T**. Similar to the derivation in section 3.1, we will again arrange the eigenvalues of **T**, i.e., the diagonal entries of **J**, in non-increasing order.

There are a few noteworthy points about factoring the matrix **T** by its generalized eigenvectors and its Jordan normal form. First, note that the generalized eigenvecotrs are linearly independent and form a basis for *n*-dimensional vectors, where *n* is the size of **T**. We denote by *n*_*k*_ the size of the Jordan block **J**_*k*_, and let 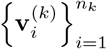 be the generalized eigenvectors corresponding to the eigenvalues in **J**_*k*_. Recalling from the notation in section 3.1, the initial distribution over the GRNs with a non-zero viability and a non-zero relative reproductivity 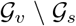 can be written as a linear combination of the generalized eigenvectors

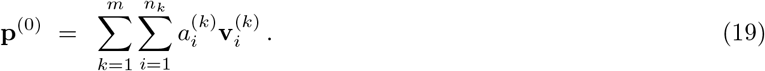

Second, since **T** is a positive matrix (see section 3.1), by the Perron-Frobenius theorem, the size of the first Jordan block *n*_1_ equals to 1. Specifically, the only entry in **J**_1_ is the leading eigenvalue *λ*_1_, and |*λ*_1_| > |*λ*_*k*_| for any *k* = 2, 3, … , *m*. For convenience, we abuse the notation and write 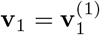.

From the matrix form of the master equation (5), we know that **p**^(*t*)^ is proportional to **T p**^(*t*−1)^, and consequently

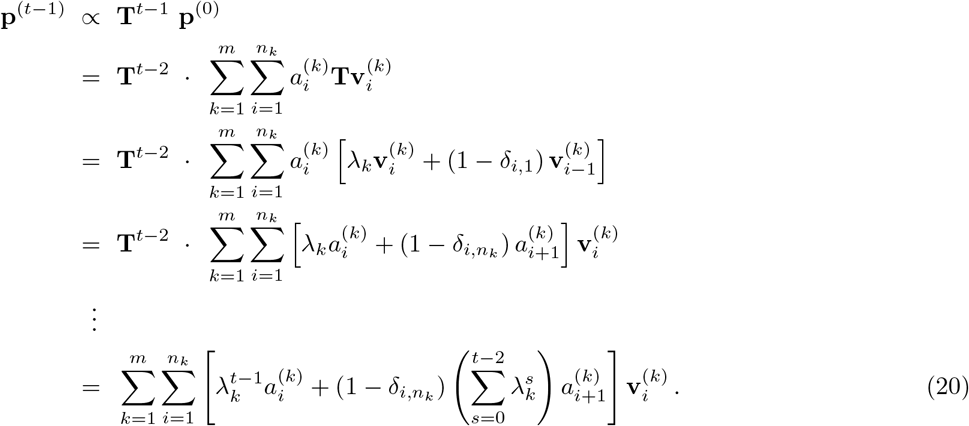

Plugging (20) into the master equation (5), we have

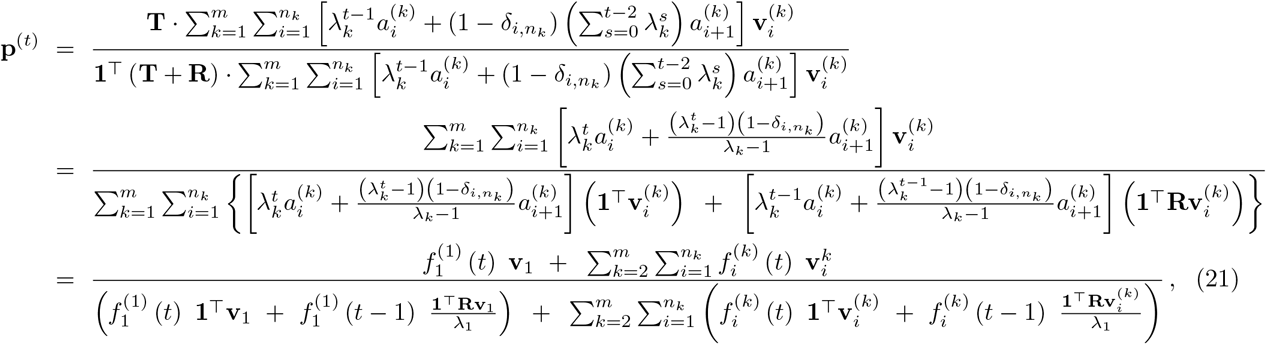

where

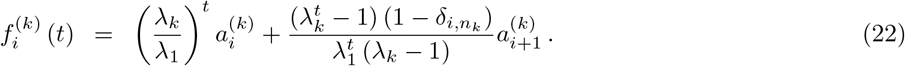

Since 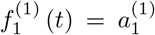 for any *t* and 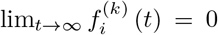 for *k* > 1, we hence recover equation (8) and show the convergence of the master equation (5) for a non-symmetric matrix **T**.

## E Convergence Rate to the Stationary Distribution of GRNs

In this section, we provide an estimate of the rate that the master equation (4) converges to its stationary distribution, using the technique known as the uniform minorization condition of Markov chains [79]. Specifically, given a sequence of probability distribution 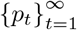 over a finite discrete space 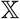 which converges to *π* = lim_*t*→∞_ *p*_*t*_, we will find an upper bound of the variation distance

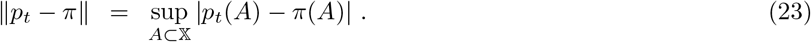

An upper bound of ║*p*_*t*_ − *π*║ will then lead us to estimating a large enough *t* such that ║*p*_*t*_ − π║ < *ϵ* for any arbitrary tolerance *ϵ*.

To begin, for any genotype/GRN *g,* 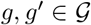, we introduce the notation

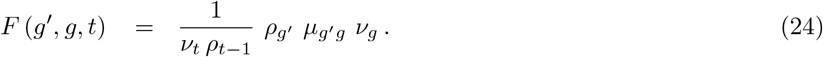

Since *v*_*t*_ = ℙ [Ψ (*I*_*t*_)] and *ρ*_*t*−1_ = ℙ [*I*_*t*−1_ → *I*_*t*_ | Ψ (*I*_*t*−1_)] are always less than or equal to 1, observe that

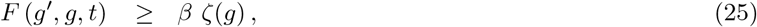

where

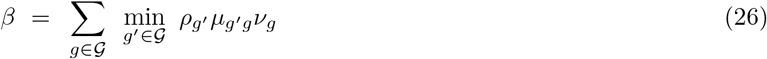

and *ζ* is a probability distribution over 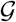 such that 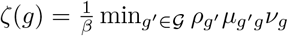 for any 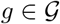. The inequality (25) is a uniform minorization condition, which has been recognized to elegantly estimate the convergence rate of Markov chains. We will adapt the derivation for Markov chains as reviewed by [79] and only summarize the key steps in what follows.

Let *X*_1_, *Y*_1_ be two independent random variables, whose probability distribution are

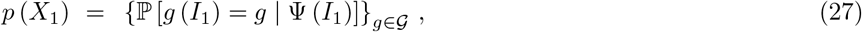

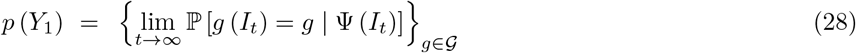

respectively. Next, given random variables *X*_*t*_ and *Y*_*t*_, let *X*_*t*+1_ and *Y*_*t*+1_ be two random variable such that

i. With probability *β*, set *X*_*t*+1_ = *Y*_*t*+1_ which follows the probability distribution *ζ*;
ii. Otherwise, *X*_*t*+1_ and *Y*_*t*+1_ are independent random variables such that

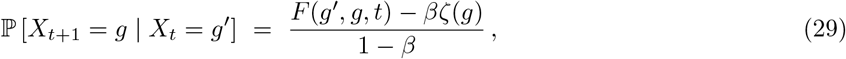

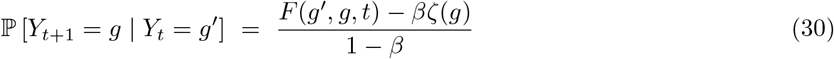

for *g,* 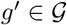.

Note that the probability distribution of 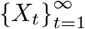 reconciles with the solution of (4) with initial condition *p* (*X*_1_), and the probability distribution of 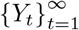 remains to be the stationary distribution of (4). We write *p*(*X*_*t*_) = 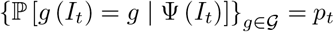 and 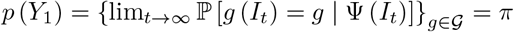 respectively.

Suppose random variable *T* to be the first time step that scenario (i) occurs so *X*_*T*_ = *Y*_*T*_. By construction, we have ℙ [*T* > *t*] = (1 − *β*)^*t*^. Let 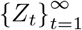 be another sequence of random variables such that *Z*_*t*_ = *Y*_*t*_ for *t* ≤ *T* and *Z*_*t*_ = *X*_*t*_ for *t* > *T*. We observe that the probability distribution of 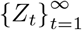 also remains to be the stationary distribution of (4). It is not hard to see that the variation distance between two probability distributions is bounded from above by the probability that the two corresponding random variables are not equal (for details, see [79]), and

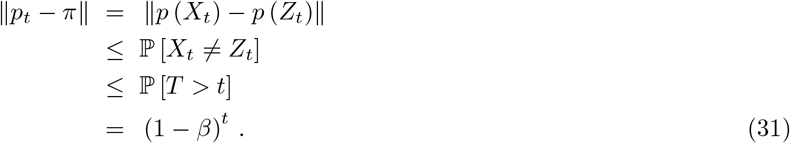

Therefore, for an arbitrary tolerance *ϵ* and in the case that there is no 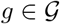 with a zero *ρ*_*g*_ (so *β* > 0), a sufficient condition for ║*p*_*t*_ − *π*║ < *ϵ* is

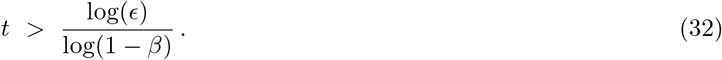

## F Supplementary Figures

**Figure 5:**
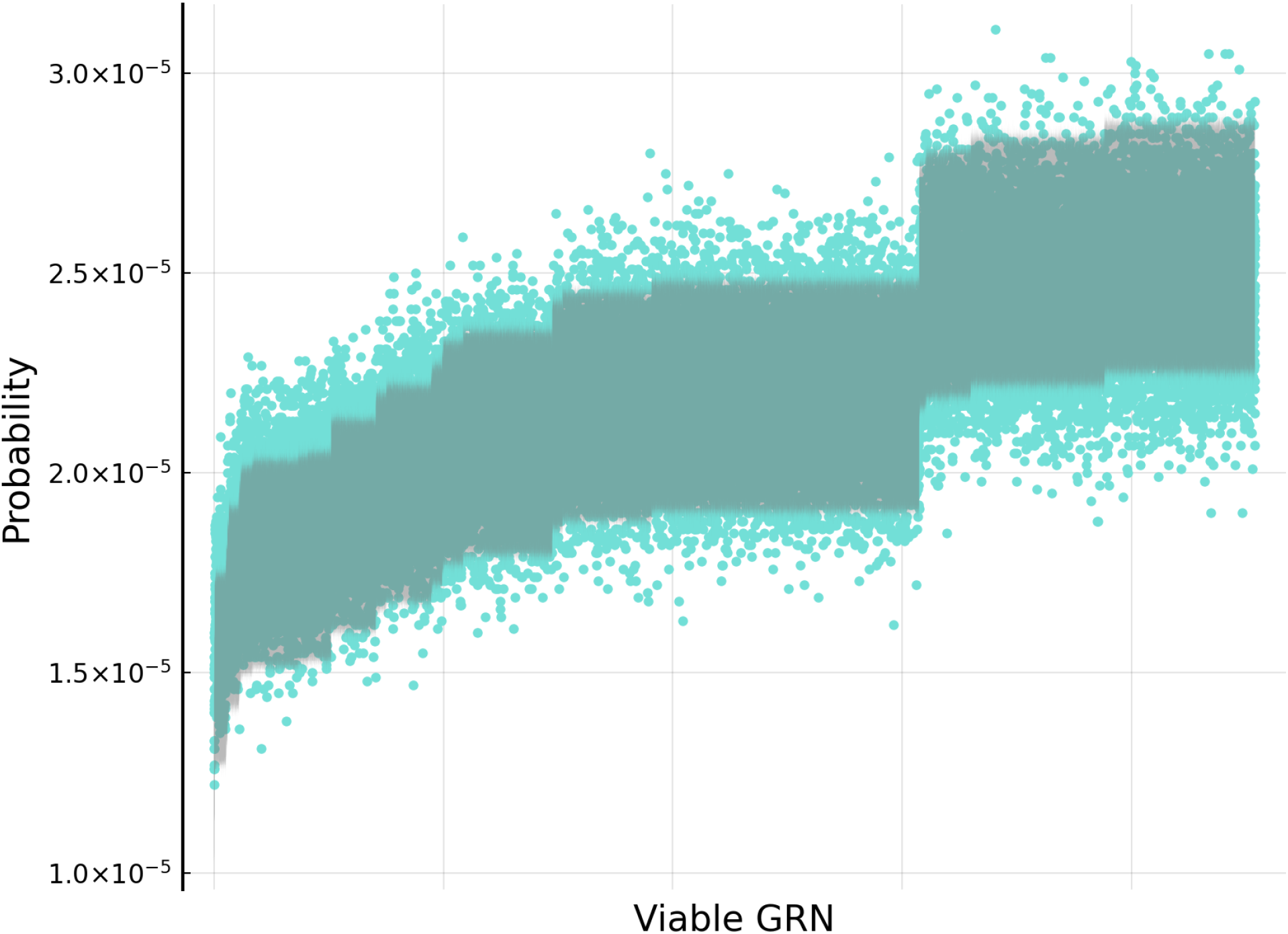
Validation that the evolutionary dynamics of GRNs converges to the derived stationary distribution. We compare the distribution of GRNs sampled from their simulated evolutionary dynamics in subsection 3.2 (blue) with the exact leading eigenvector of the transition matrix (11) with the same *μ* = 0.1 (gray). The shaded area shows its 95% confidence band that accounts for the uncertainty of finite-sized sampling in the simulations. No significant deviation was found.

## Footnotes

1 A set *A* is said to be *partitioned* by 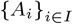 if *A* = ⋃_*i*ϵ*I*_ *A*_*i*_ and *A*_*i*_ ∩ *A*_*j*_ = ∅ for two distinct *i, j* ϵ *I*.

2 To be more precise, this argument is only valid when the joint probability for any combination of multiple single-locus mutations is non-zero per generation. Otherwise, we can modify the master equation (5) by extending the time scale from 1 to Δ*t*, where Δ*t* is the diameter of the subgraph of the genotype network constrained by a non-zero viability and reproductivity. The modified transition matrix is now proportional to **T**^Δ*t*^, which is a positive matrix since mutation events at different generations are independent. Replacing **T** by **T**^Δ*t*^ we have an analogous derivation to prove the convergence of the master equation.

3 Here we abuse the notation v_1_ (**A**) and *λ*_1_ (**A**) for the leading eigenvector and the leading eigenvalue of **A** respectively.

4 We denote by 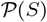 the *power set* of a set *S*, which is the set of all possible subsets of *S*.

5 Here such a rewiring process through an intermediate gene is necessary for most scenarios. Specifically, in the case where 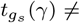 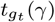, directly rewiring 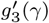 to 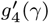 may break the essential pathway to 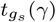. One can alternatively find a gene *γ*′ whose allele does not produce 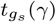 or 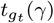 in 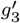, where the existence of *γ*′ is guaranteed since |Γ| > |Ω_+_|. Rewiring 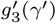 to 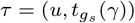 for some *u* ϵ Ω_0_, applying the direct rewiring between 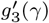 and 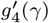, and then rewiring *τ* back to 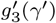 avoids the potential break of the essential pathways.

